# Decoding Study-Independent Mind-Wandering from EEG using Convolutional Neural Networks

**DOI:** 10.1101/2020.12.08.416040

**Authors:** Christina Yi Jin, Jelmer P. Borst, Marieke K. van Vugt

## Abstract

**Objective:** Mind-wandering is a mental phenomenon where the internal thought process disengages from the external environment periodically. In the current study, we trained EEG classifiers using convolutional neural networks (CNN) to track mind-wandering across studies.

**Approach:** We transformed the input from raw EEG to band-frequency information (power), singletrial ERP (stERP) patterns, and connectivity matrices between channels (based on inter-site phase clustering, ISPC). We trained CNN models for each input type from each EEG channel as the input model for the meta-learner. To verify the generalizability, we used leave-N-participant-out crossvalidations (N=6) and tested the meta-learner on the data from an independent study for across-study predictions.

**Main results:** The current results show limited generalizability across participants and tasks. Nevertheless, our meta-learned trained with the stERPs performed the best among the state-of-theart neural networks. The mapping of each input model to the output of the meta-learner indicates the importance of each EEG channel.

**Significance:** Our study makes the first attempt to train study-independent mind-wandering classifiers. The results indicate that this remains challenging. The current proposed stacking neural network design allows an easy inspection of channel importance and feature maps.

## 1 Introduction

Mind-wandering is a thought process that is characterized by being not directly relevant to the primary goals in the current context (Smallwood & Schooler, 2015). Mind-wandering tends to manifest itself as attentional lapses which often contribute to making errors in a task (Cheyne et al., 2006). However, mind-wandering does not result in errors in all cases. Sometimes people can handle their primary tasks well when they start enjoying these periods of self-distraction as a temporary escape from the current situation (Schooler et al., 2011). This positive effect of mind-wandering is particularly common when the current task has a low cognitive load—in other words, performance can be achieved automatically with little executive control involved (Randall et al., 2019) and therefore low performance is not a foolproof indicator of mind-wandering. Another behavioral measure that has been proposed to characterize mind-wandering is increased response time variability. Several studies have shown that even when no obvious mistakes are observed, participants display increased variance in their response times when their mind wanders (Bastian & Sackur, 2013; Seli et al., 2013; Zanesco et al, 2021b; Zheng et al., 2019).

In addition to using behavior, mind-wandering can also be detected by means of physiological and neural measures. For example, researchers found that the pupil diameter in reaction to stimuli becomes smaller in an off-task state (Huijser et al., 2018), possibly related to a vigilance decrement that tends to co-occur with mind-wandering (Unsworth & Robison, 2016). On the level of the cerebral cortex, mind-wandering appears to be associated with inhibited sensory processing to the visual stimuli, referred to as ‘perceptual decoupling’ (Schooler et al., 2011). This perceptual decoupling manifests itself in the EEG as a reduced P1 and increased alpha power (frequency range 8.5 ∼ 12Hz) observed at the parietal-occipital regions (Compton et al., 2019; Jin et al., 2019; Kam & Handy, 2013). In fMRI studies, mind-wandering is associated with increased activation of the default mode network (DMN), together with changes in the connectivity between the DMN and other networks (Christoff et al., 2009; Ho et al., 2019). Through indicating a memory retrieval process, the involvement of the DMN supports the functional role of mind-wandering as “spontaneous future cognition” (Cole & Kvavilashvili, 2019), potentially aiding problem-solving and creativity (Schooler et al., 2011).

Given this set of neural and physiological correlates of mind-wandering, several studies have explored the possibility of predicting mind-wandering on a single-trial level using machine learning. Mittner and colleagues used multimodal signatures of the (co)activations in the DMN, ACN (anti-correlated network), and the pupil diameter as the features for a machine learning model to predict mind wandering (Mittner et al., 2014). They found that neural data could reliably predict mind-wandering with a median accuracy of 79.7% using leave-one-participant-out cross-validation (LOPOCV). They also found that mind-wandering was not linked to DMN, CAN or pupil diameter alone, but that instead all the features were necessary for the optimal predictive performance (Mittner et al., 2014). This seems sensible, given that the contents of mind-wandering can vary considerably. In their recent work, they achieved 65% accuracy *across* participants by training another multimodality machine learning model (Groot et al., 2021). The authors attributed the different performance level achieved across studies to individual biases in self-reports, differences in levels of meta-awareness, and heterogeneity in the thought content being reported (Groot et al., 2021).

Kawashima and Kumano (2017) found that training an EEG-based mind-wandering classifier with a non-linear support vector machine (SVM) performed better than a linear SVM. They also found that training with a selected subset of electrodes from frontal and parietal-occipital regions performs better than using all the electrodes. This suggests that the correlation between EEG and mental state fluctuations is more complex than a simple linear relationship and a subset of electrodes at key positions might contain sufficient information for discriminating mind-wandering. Jin et al. (2019) explored whether it was possible to predict mind wandering on the basis of various features of EEG data, ranging from power and inter-site phase clustering in the alpha and theta bands, to single-trial ERPs. They were able to predict mind-wandering with an average accuracy of 64% for a sustained attention to response task, and 69% for a visual search task. Additionally, they performed across-task predictions between the sustained attention to response and visual search tasks, with an accuracy of 59% and 60% (Jin et al., 2019). In a related study, the researchers used ICA-decomposed EEG in the alpha band to predict the occurrence of mind-wandering—albeit this time in a model that generalized across participants. They reported again an average accuracy of around 60% when testing the models on a left-out dataset (LOPOCV), for both within- and across-task predictions (Jin et al., 2020).

Several conclusions can be drawn from the studies above. The accuracy with EEG seemed to hit a ceiling at 60% when making generalization predictions across individuals or across tasks. One possibility is the relatively low signal-noise ratio (SNR) of scalp EEG compared to other neural imaging techniques, such as intracranial EEG or fMRI. However, scalp EEG is still one of the main signals to be used for Brain-Computer Interfaces with healthy users. Another possible cause for low accuracy is that labels come from thought probes. Other setups, for example facial video recording, eye-tracking measures and pupillometry can potentially help to validate and revise the behavioral scales. A third possible cause is that learning is performed with pre-computed EEG features (e.g., P3 or alpha power). These features are selected based on previous studies, but they do not represent all the temporal-frequency information from EEG sensors across the whole scalp. Also, the studies discussed above trained SVMs to learn the relationship between pre-computed features. SVM—when used with a nonlinear kernel—is a powerful tool for learning features following non-linear relationships, but its computational cost is relatively high (O(n^3^), Abdiansah & Wardoyo, 2015). To handle a large quantity of the frequency information and/or evoked potentials from all the sensors, we need more powerful machine learning models.

The convolutional neural network (CNN) is a good candidate for addressing this type of problem. A CNN uses a kernel to detect the features of the input signals. With deeper CNN layers, the learned features become more abstract, while at the same time the dimensions of the input decrease. The CNN layers use much fewer parameters than the fully connected (FC) neural networks to reduce the computational and storage cost. In practice, multiple CNN layers are designed to detect useful features. Then the fully connected layers learn the relationship of the features for the final prediction. Hosseini and Guo (2019) trained a CNN architecture to detect mind-wandering in a meditation task^1^. They trained datasets from two participants and achieved the highest performance of 91.78% during 10fold cross-validation (CV) within-individuals. The authors also tried to predict across individuals by training a classifier with each individual only and using it to predict the other dataset. The performance dropped to 66% for across-individual predictions. Their results indicate that the generalizability of mind-wandering classifiers is challenging, even with state-of-the-art neural networks.

Nevertheless, Hosseini and Guo (2019) only used the raw EEG signal to train their CNN classifiers, which limits the performance. In addition, they had a very limited sample size. In the current study, we will include band-frequency information and stERPs as input types for the CNN models as well as the raw EEG. Furthermore, we endeavor to train study-independent mind-wandering classifiers: we trained and validated classifiers with data from Jin et al. (2019, referred to as Study A) and tested the classifiers on the data from Jin et al. (2020, referred to as Study B). In addition, each study contained two independent tasks and different groups of participants participated in each of these studies. The design and the probes of both studies can be found in Figure 1.

**Figure 1.**
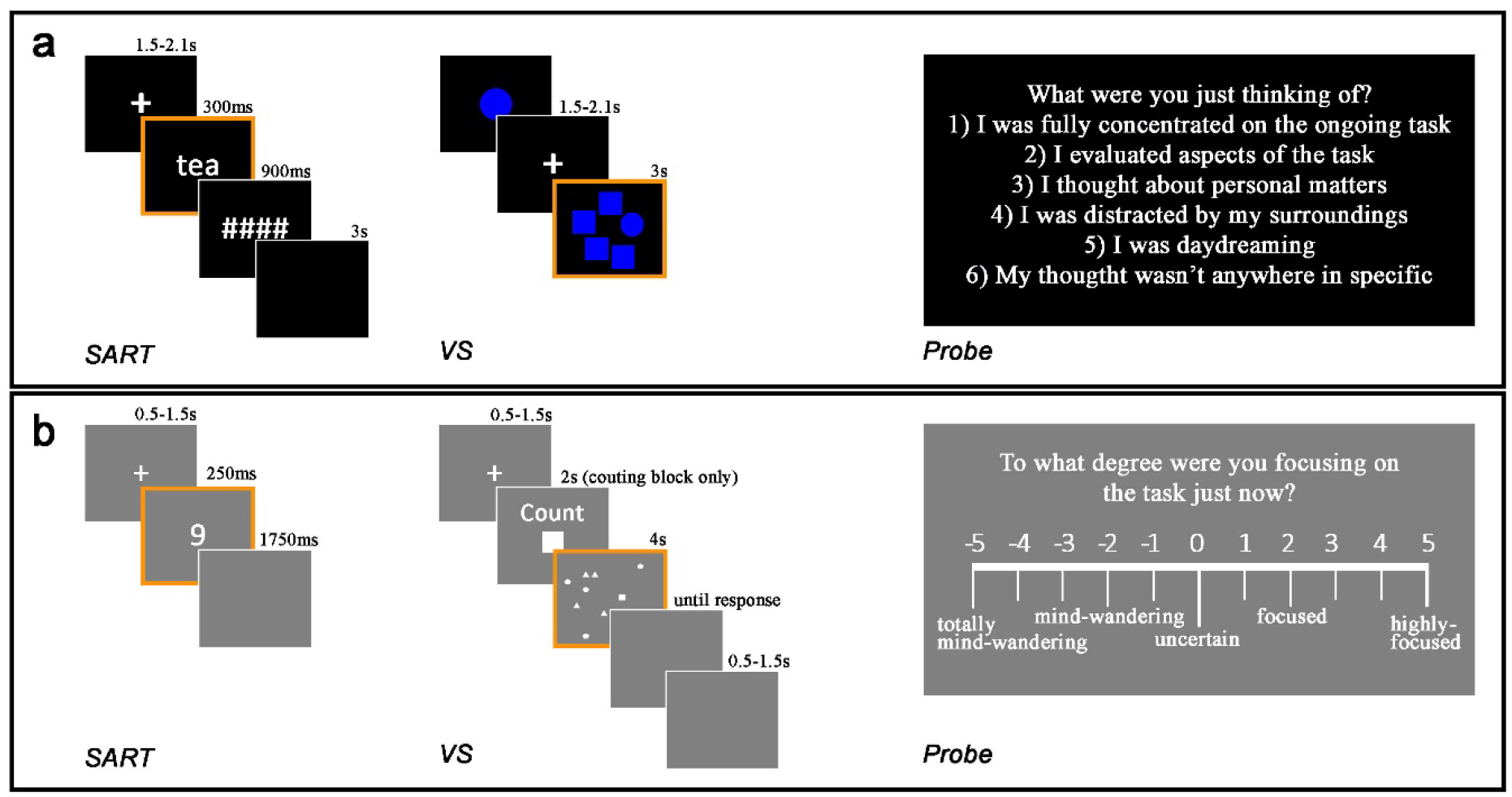
Trial sequences of the SART and the visual search (VS) task, and the probe illustrations for the (a) training and (b) testing dataset. The orange frame indicates the stimuli around which the EEG epochs were extracted (each epoch comprised the period starting at stimulus onset).

For developing the CCN model, we first transformed the raw EEG into the frequency power spectrum and stERP contour maps. We also included the inter-site phase clustering (ISPC) reflecting connectivity between EEG channels. Raw EEG, power and stERP from each channel and the ISPC from each channel pair was trained with an independent CNN classifier. The output for each model represents the activations of one on-task neuron and one mind-wandering neuron using one input type from one spatial point (channel or channel pair). To figure out the relationship between the input types and their spatial sampling points, we trained a meta-learner with the concatenated binary outcomes of each input model (Figure 2). This allowed us to evaluate the contribution of each input model by mapping their weights to the final output layers of the meta-learner. To compare the performance of our classifier to the other state-of-the-art neural networks, we trained two other CNN models – the beforementioned Hosseini and Guo (2019) model and EEGnet (Lawhern et al., 2018). The EEGnet is superior in learning frequency patterns with only raw EEG as the input. Its architecture design also allows to map the feature contribution to its spatial patterns. The EEGnet has been proven effective at solving classic BCI problems such as motor imagery and P300 speller (Lawhern et al., 2018).

**Figure 2.**
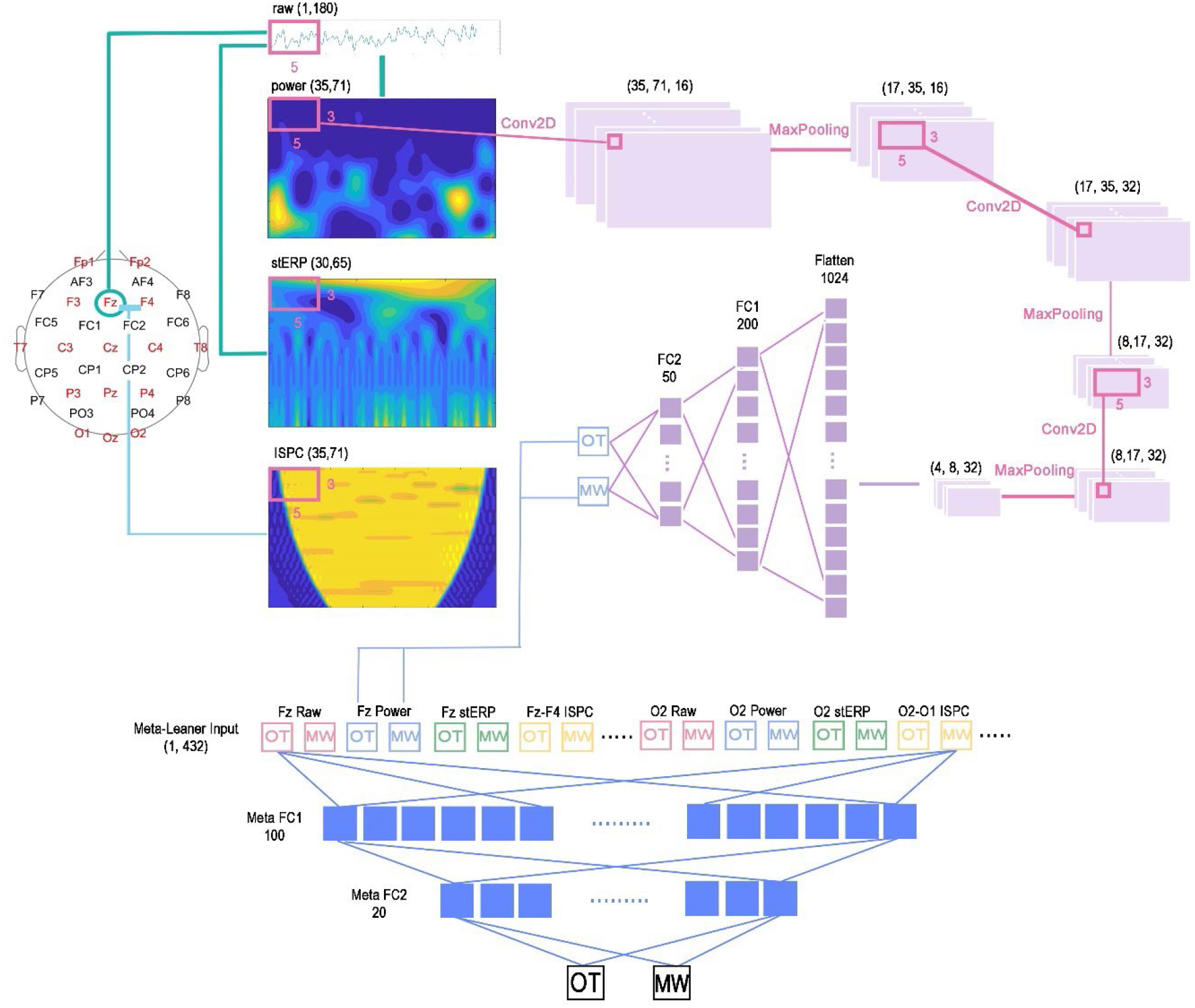
Channel locations (32-channel 10-10 system). EEG from each channel is transformed into power spectrum and single-trial ERP (stERP) contour maps. The 16 channels in red font were also used to compute the inter-site phase clustering (ISPC). Each input type from each spatial location (channel or channel pair) was trained with three convolutional layers (of which neuron numbers are 16, 32 and 32) with a max-pooling layer after each CNN. For raw EEG, we used a 1D CNN with a kernel length of 5 to account for the pattern of every five temporal sampling points. For other types of input, we used a 2D CNN with a kernel size of (5,3) to learn the pattern of 5 temporal sampling points and 3 frequency sampling points. The obtained model with each input type from one spatial location serves as the input model for the meta-learner, which learned the relationship of the binary outcomes between each input model with two fully connected (FC) layers and generated the final prediction.

## 2 Method

### 2.1 Datasets

#### 2.1.1 Training dataset

The training dataset is derived from Jin et al. (2019). Thirty participants (13 females, ages 18–30 years, M= 23.33, SD = 2.81) took part in the study. They performed a visual search task and a sustained-attention to respond task (SART) for six blocks each in two sessions (Figure 1a). The main stimuli in the SART were English words that occurred in lowercase for 89% of the time and in uppercase for 11% of the time. Participants were required to press “m” whenever they saw a lowercase word and to withhold their response when an uppercase word appeared. In the visual search task, participants were given a target shape to search at the beginning of each block. They were required to look for the target in each trial of that block and indicated if the target was in the search panel (yes/no) by pressing the left or right arrow key corresponding to their response. There was an equal probability of the target-present and the target-absent trials. A SART block had 135 trials and a visual search block had 140 trials. Details can be found in the original study (Jin et al., 2019).

Participants were interrupted by probe questions asking them about the content of their thinking at that moment (Figure 1a). They could respond to this question with one of six options: (1) I was entirely concentrated on the ongoing task; (2) I evaluated aspects of the task (e.g., my performance or how long it takes); (3) I thought about personal matters; (4) I was distracted by my surroundings (e.g., noise, temperature, my physical condition); (5) I was daydreaming, thinking of task-unrelated things; (6) I was not paying attention, but my thought wasn’t anywhere specifically. Each task had 54 probes that were interspersed with the trials. Two consecutive probes were separated by 7–24 trials, roughly accounting for 34–144 seconds. In the analysis, response 1 and 2 were labelled as an on-task state; response 3 and 5 were labelled as a mind-wandering state, and the other responses were ignored, following the same classification rule as the original work.

#### 2.1.2 Testing dataset

The testing dataset is derived from Jin et al. (2020). Thirty participants (16 females, age 18–31 years, M = 23.73, SD = 3.47) took part in the study (Figure 1b). In the visual search task, participants either counted the specified target in the following search panel and indicated their response by pressing the number key (counting condition), or passively viewed the search panel and pressed ‘j’ as a standard response (non-counting condition). In the SART, participants viewed single digits ranging between 1 to 9 drawn from a uniform distribution. They pressed ‘j’ whenever they saw a digit other than ‘3’; if ‘3’ appeared (11%), they were to withhold their response. The visual search task had 21 trials in each block and 20 blocks in total. The SART had twelve blocks with a block length that varied between two to seven repetitions of the nine digits (18-63 trials). Probes were shown at the end of each block in both tasks. Block length varied between one to three minutes. Further details can be found in the original paper (Jin et al., 2020).

In this dataset, the probes appeared at the end of each block (which were much shorter than in the training dataset). Participants indicated their momentary attentional state on a rating scale of -5 to 5 with anchor ‘-5’ for ‘totally mind-wandering’, ‘-2’ for ‘mind-wandering’, ‘0’ for ‘uncertain’, ‘2’ for ‘focused’, and ‘5’ for ‘highly focused’. Thus, positive ratings were classified as on-task and negative ratings as mind-wandering.

### 2.2 EEG preparation

#### 2.2.1 Recording hardware

The training dataset had 128 channels, while the testing dataset had 32 channels. All electrode locations were within the International 10-10 System. In the current analysis, we only considered the 32 channels that overlapped between the studies (Figure 2). The data were recorded using the Biosemi ActiveTwo recording system. The online sampling rate was 512Hz. The Biosemi hardware does not have any high-pass filtering. An anti-aliasing filtering is performed in the ADC’s decimation filter (https://www.biosemi.com/faq/adjust_filter.htm).

#### 2.2.2 Preprocessing

Data had already been preprocessed in the original studies, and we reused these preprocessed data here. The offline EEG preprocessing was done with the EEGLAB toolbox in MATLAB. Continuous EEG was re-referenced to the average signal of both mastoids. The band-bass filtering was set to be 0.5-40 Hz and 0.1-42 Hz for the training and testing datasets, respectively. Both datasets were downsampled to 256 Hz. The original segmentation was [-400 1200] ms with respect to stimulus onset for Study A, and [-1000 3000] ms for Study B. In the current study, we took the overlapping [-400 1000] ms as the time window for analysis. The 200ms before the stimulus onset was used as the baseline. Ocular artifacts were detected and removed using the infomax independent component analysis (ICA).

#### 2.2.3 Transformation

Apart from the raw EEG data, three kinds of transformations were performed on the original EEG matrix – power and inter-site phase clustering derived from a time-frequency decomposition using the complex Morlet wavelets (Cohen, 2014), and single-trial ERP (stERP) analysis (Bostanov, 2004; Bostanov & Kotchoubey, 2006; Jin et al., 2019). These transformations were performed in MATLAB with customized code.

A complex Morlet wavelet is created using the equation

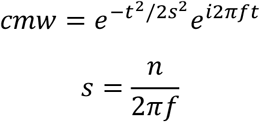

where f denotes the frequency in Hz, and n refers to the number of wavelet cycles. Convolving the complex wavelet with the original signal gives complex dot products, from which the amplitude can be extracted through the vector length of each complex data point (power is obtained from the squared amplitudes, Figure 2c, and the phase angle is obtained from the angle with respect to the positive real axis).

A set of wavelets was created with frequencies ranging from 4 to 40 Hz with 35 frequency sampling points in the logarithmic space, covering multiple frequencies in each of the theta/alpha/beta/gamma bands. Delta oscillations (1-3Hz) were excluded because our time window of 1400ms was not long enough to estimate delta power with sufficient accuracy. The upper boundary of the analyzed frequency was limited by the bandpass filtering during the preprocessing. The number of cycles used for the wavelet increased from 3 to 7 in a logarithmic spacing on the frequency axis.

Inter-site phase clustering (ISPC) was computed as a measure of the connectivity between channels (Cohen 2014):

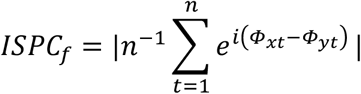

in which Φ_x_ and Φ_y_ are the phase angles from electrode x and y. ISPC is computed as the averaged phase angle differences (which were also mapped to the complex plane) in a moving time window. The length of the averaged angle difference denotes the clustering (i.e., the more the phase difference remains constant, the more their length adds up). The window length increased from 3 to 5 cycles linearly with the frequency. As shown in Figure 2, there is an empty region of half the wave length at both sides of each frequency with no averaging data. Those regions were set to be zero during machine learning. The ISPC was computed in a 16-channel layout resulting in 120 channel pairs (Channel layout in red font, Figure 2) to reduce the number of channel pairs.

Both the power and ISPC matrices were down-sampled to 50Hz given the fact that time-frequency analysis smears out the signal over time.

The single-trial ERP analysis, which quantifies the bumps in the EEG in terms of temporal location and amplitude was performed through computing the cross-covariance between the signal and the kernel *ψ*(t):

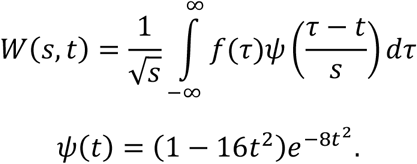

With two varying parameters - time lag τ and scale s (an indication of wavelength), the cross-covariances can be mapped to a contour graph (Figure 2e). In the current study, the time lag (ττ) ranges from 0 ms (stimulus onset) to 1000 ms (end of the EEG epoch) following the same temporal resolution as the raw EEG. The scale (s) ranges from 1 to 2500 ms in logarithmic space with 300 sampling points. The obtained matrix is down-sampled to 65 sampling points in the time lag and 30 points in the scale (frequency dimension).

Together, this creates four types of inputs for the CNNs: raw EEG, power, ISPC, and stERP (Figure 2).

### 2.3 Neural Network

#### 2.3.1 Architecture of the neural network

The CNN input model and the meta-learner structure are shown in Figure 2. For the raw EEG input, we used three layers of a 1D CNN with a kernel size of five temporal points. The neuron numbers are 16, 32 and 32 sequentially^2^. Each convolutional layer is followed by a max-pooling layer with a pooling size of 2. The output of the third max-pooling layer is flattened and connected to two FC layers with 200 and 50 neurons. The activation function is ReLU for all the hidden layers. The output layer uses a softmax activation function to calculate the activation of one on-task neuron and one mind-wandering neuron. The CNN structure for the power, ISPC or stERP is similar to that of the raw EEG except that a 2D CNN replaced the 1D CNN. The kernel size of the 2D CNN is (3,5) to account for the pattern of three frequency points and five temporal points. Each 2D CNN layer is followed by a max-pooling layer with a pooling size of (2,2). The learning was performed with categorical cross-entropy as the loss function and optimized by Adam at a learning rate of 0.0001. The training batch size is 120, iterated over 200 epochs.

The meta-learner is trained with the concatenated binary outcomes from each input model. For the raw EEG, power and stERP, we trained 32 input models (using data from each channel). For the ISPC, we trained 120 input models (using phase clustering from each channel pair). Thus, altogether, we had 2×32×3+2×120=432 outputs from all the input models to form a (1, 432) vector as the input to the meta-learner (Meta_Full). We also trained two other meta-learners using a subset of those input models. One is without the power but with the other three input types (Meta_NoPower), because power was the most overfitting input according to a preliminary analysis (suppl. II). The third metalearner used only stERP (Meta_stERP), as this input type performed best during validations in the preliminary study (suppl. II).

The meta-learner used two FC layers consisting of 100 and 20 neurons. The final output consisted of two neurons that indicate the prediction of on-task or mind-wandering. The hidden layers of the metalearner used standard ReLU activation functions and the output layer used softmax. The meta-learner was trained with batches of 200 trials and iterated over 50 epochs, performed by categorical crossentropy as the loss function and optimized by Adam at a learning rate of 0.0005.

We set the dropout to be 0.4 to train the input models and 0.2 to train the meta-learner in order to prevent overfitting. The proposed CNN models and meta-learner is implemented on a workstation with an Intel Xeon 2.2 GHz, 512GB RAM, and a TITAN X, TITAN Xp, and two GeForce RTX 2080 graphical cards with CUDA V10.1.243 (one or two of the four GPUs depending on the availability) using Python 3.7 and the keras machine learning library.

#### 2.3.2 Validation and Testing

A leave-N-participant-out cross-validation (LNPOCV) was used to assess the performance. The N was decided to be 6 to account for 20% of the datasets given that the total sample consisted of 30 participants. In that sense, we are performing 5-fold CV across individuals. In each fold, we set aside 6 individual datasets for validation purposes and trained the classifiers with the other 24 individual datasets. Each individual dataset was used once for the validation. The performance was indicated by both the accuracy and the area under the curve (AUC) of the Receiver Operating Characteristic (ROC).

#### 2.3.3 Comparison models

We trained a CNN using the architecture by Hosseini and Guo (2019)^3^ and an EEGnet^4^ for comparison purposes. Both used only raw EEG from multiple channels as input. The validation and testing procedures remained the same.

## 3 Result

### 3.1 Network performance

We used trials within 12 seconds before each probe for both the training and testing samples. This time window results in 3169 on-task (OT) trials and 1976 mind-wandering (MW) trials from Study A, and 1266 OT and 424 MW trials from Study B.

We first examined the best normalization approach for each input type in a preliminary analysis. Given that the classes were not balanced originally, we adjusted the weight of the MW class while keeping the weight of the OT constant at 1. The best normalization and class weights for each input type can be found in Suppl. I and II. Input types were normalized with the best normalization and trained with the best class weights to be the input models for the meta-learner.

The performance of the meta-learners, as well as the comparison models, are listed in Table 1. The models learned well during the training, with above 75% of the AUC for all models. However, the classification performance dropped when predicting the left-out datasets. The meta-learner with stERP performed relatively the best during validations, achieving 59% accuracy and .57 for the AUC, indicating a mild level of generalizability across individuals. The meta-learner with power as the input (Meta_Full) and the Hosseini2019 model showed strong signs of overfitting. They both correctly identify almost 100% of the training labels. However, their performance on the validation datasets was almost at chance level (ROCAUC .505 and .509 during LNPOCV).

**Table 1.** Network performance as indicated by accuracy and ROC area under the curve (AUC). The Three meta models indicate the meta-learner with all the input types (Meta_Full), without power for input (Meta_Power) and with single-trial ERP only (Meta_stERP).

|  |  | Meta_Full | Meta_noPower | Meta_stERP | Hosseini2019 | EEGnet2018 |
| --- | --- | --- | --- | --- | --- | --- |
| Training | Accuracy | .980 | .828 | **.806** | 1 | .754 |
|  | ROCAUC | .980 | .828 | **.806** | 1 | .754 |
| Validation<br>(LNPOCV) | Accuracy | .534 | .579 | **.587** | .539 | .468 |
|  | ROCAUC | .505 | .565 | **.569** | .509 | .450 |
| Testing | Accuracy | .511 | .492 | **.519** | .545 | .457 |
|  | ROCAUC | .501 | .485 | **.506** | .492 | .539 |

Finally, we tested all the models by predicting the data of Study B. The performance decreased further, as expected. Only two of our meta-learners (Meta_Full and Meta_stERP) can predict above chance level in both accuracy and AUC. The meta-learner with stERP (accuracy .519 and .506) is slightly better than the meta-learner with all input types. Interestingly, Hosseini2019 achieved the best accuracy (.545) during the testing while the ROCAUC was .492. On the contrary, EEGnet2018 had the highest ROCAUC (.539) but the accuracy was the lowest (.457). Our Meta_stERP model achieved balanced performance in both accuracy and ROCAUC.

### 3.2 Learned features

We investigated the importance of the input locations by analyzing the weights of the input model in the meta-learner. Given that Meta_stERP performed the best, we mapped location importance using weights derived from this model. The output of the meta-learner consisted of one neuron responding to the OT state and another neuron responding to the MW state. As each input model also represented one OT neuron and one MW neuron, the mapping gives two weight topoplots for the meta-OT neuron (OT-OT, MW-OT) and two weight topoplots for the meta-MW neuron (OT-MW, MW-MW). To simplify the interpretation, we performed PCA on the two weight topoplots for each meta-output neuron and obtained the largest PC accounting for 100% of the variance (Figure 3A). The importance of channel locations in the PC topoplots looks similar between the meta-OT and the meta-MW neurons. Some channels are important in both OT and MW activations (e.g., P4). Some other channels are responded to more by one of the two classes (e.g., C3 decides the activations of the meta-OT neuron more than the meta-MW. O2 decides the activation of the meta-MW neuron more than the meta-OT).

**Figure 3.**
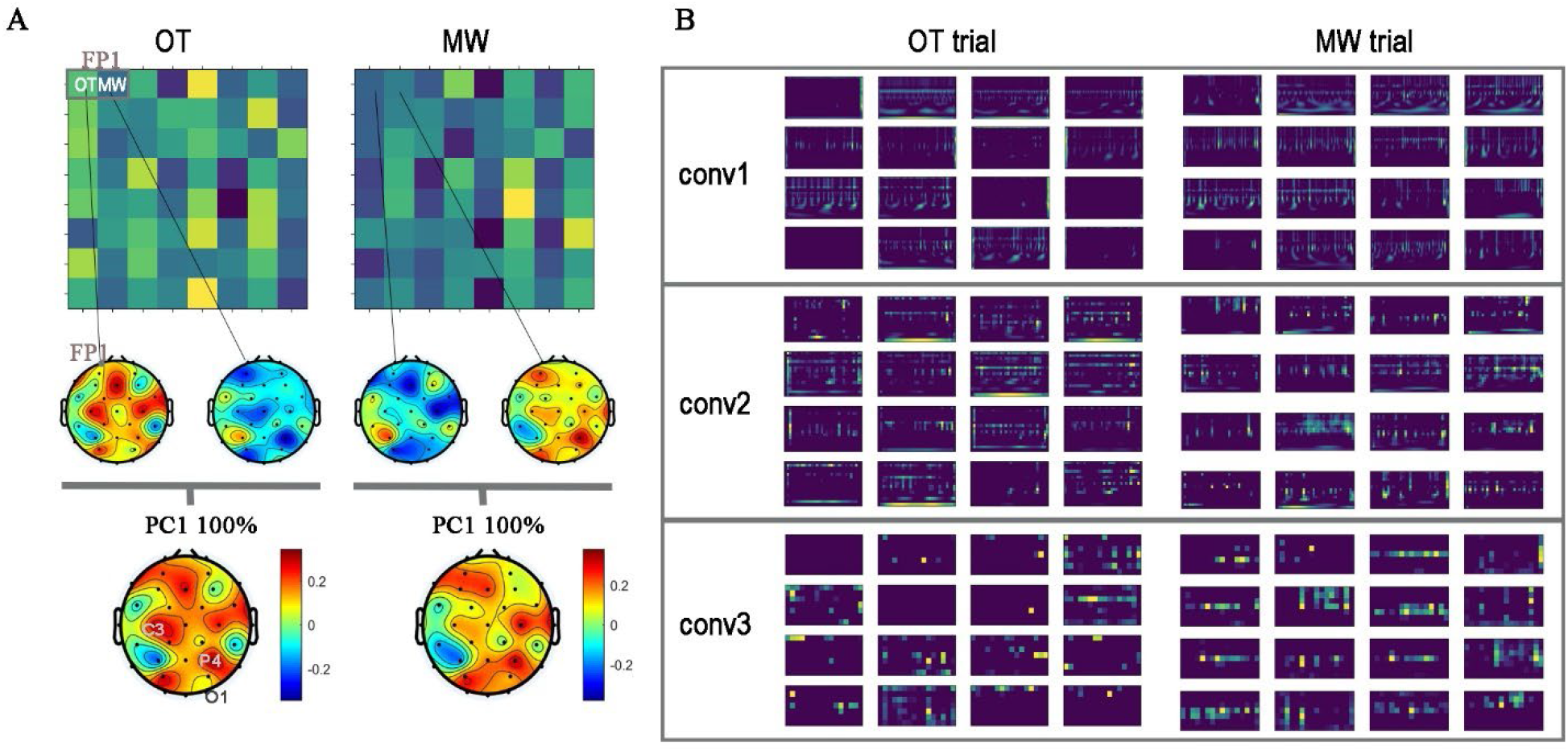
(A) Mapping of channel importance of the meta-learner trained with stERP. The weights of each input model to the meta-learner outputs are marked in the matrices. Each matrix is further mapped to two weight topoplots for the input of OT and MW activation, respectively. To simplify the interpretation, the weight topoplots are decomposed by the PCA to obtain the largest PC for each meta-output neuron. (B) Feature maps of each convolutional layer with one OT and MW trial as examples.

We chose P4 as an example to plot the feature maps because it is informative in activating both the meta-OT and meta-MW neurons (Figure 3A). Feature maps are based on the output of each convolutional layer (Figure 3B). Comparing the feature map of one OT trial and one MW trial, especially the feature map of the third convolutional layer, we found the features are identified during the whole time series and across the whole scale range of the stERP, indicating that all the ERP components are likely to be predictive. The activation of the middle scale range seems to be highlighted in the MW trial compared to the OT trial. Given that the middle scale range is equivalent to the wavelength range looking for early sensory evoked potentials, this indicates that early sensory processing is likely to feature mind-wandering.

## 4 Discussion

In the current study, we trained meta-learning neural networks with multiple CNN input models to classify mind-wandering. Each input model was trained with one EEG input type from one spatial sampling point to predict mind-wandering separately. Thus, the meta-learner not only learned a synthesized prediction based on all the input models but also allowed us to map the importance of each channel by examining the weights of the input models to the meta-learner outputs.

The current results demonstrate the difficulty of obtaining mind-wandering detection that is generalizable across individuals or studies. The generalization across individuals is shown during the LNPOCV. While the CNN classifier and the meta-learners performed well on the training datasets (above .75 for the ROCAUC), the generalizability dropped below .6 in the accuracy and AUC. Furthermore, the performance on the testing datasets indicated the limitation of such across-study predictions.

Nevertheless, the currently proposed meta-learner with stERP as the input achieved the best performance during the validation and testing stages. We examined the channel importance by mapping the weights of each input model to the outcomes of the meta-learner and found a similar channel importance map between the final OT and MW neuron as learned by the meta-learner. Finally, by looking at the feature maps between classes, we can understand how the CNN learned the patterns from the stERP contour maps.

The largest limitation of the current study is the generalizability of the CNN classifiers, even though we addressed the problem with state-of-the-art neural networks. We attribute the main cause to the heterogeneity of mind-wandering: on one side, individuals differ in their mind-wandering thoughts as well as the patterns of the neural activation of the heterogenous thoughts (Christoff et al., 2016; Wang et al., 2018; Zanesco et al., 2021a); on the other side, mind-wandering and on-task is a dual-tasking process – individuals are likely to keep working on the primary job without the performance being interrupted if the primary task is low-demanding or habitual (van Vugt et al., 2015). In that case,“free” cognitive resources are available to be used by mind-wandering (Taatgen et al., 2021), therefore making mind-wandering generation becomes “hidden” and hard to be discriminated. Based on the current results, we recommend considering individual classifiers instead of inter-individual classifiers to detect mind-wandering for real-life BCI applications.

## 5 Conclusion

The current study indicates that a generalizable classifier to detect study-independent mindwandering episodes with scalp EEG remains challenging. Nevertheless, we found that the metalearner with input models trained with single-trial ERP contour maps performed the best. We also showed how this work can contribute to explainable artificial intelligence by giving an example of how channel contributions and the learned features can be examined by means of the weights of the input models and the feature maps.

## Data and Code Availability Statement

The data that support the findings of this study are openly available in https://unishare.nl/index.php/s/T94LXPQqw5FEA4J. Analysis code are available in https://github.com/christina109/MW_EEG_CNN.

## Footnotes

1 Mind-wandering in meditation tasks is defined as decoupling from internal sensations (e.g., focusing on the breath) instead of decoupling from processing external stimuli in a cognitive task situation such as discussed in the current paper, which most likely requires a different classifier.

2 The choice of neuron number is decided according to preliminary results, in which we tested CNN design with [16, 16, 8], [16, 32, 32] and [64, 64, 32] neurons. The results (Suppl. III) indicate an improvement on ROCAUC during validations by increasing neurons from [16,16,8] to [16,32,32] (.513, .519 respectively). However, adding more neurons to make a design as [64,64,32] did not increase the ROCAUC on validations (.519). Therefore, we decided [16,32,32] to be the proposed network design following Occam’s Razor Principle.

3 The proposed architecture accommodates input size (64, 8192), while the current data size is (32, 180). We thus reduced the kernel length to accommodate for our data size. The architecture and other hyperparameters are the same as in Hosseini and Guo (2019)

