## Supplemental Table 1 for "Decoding Study-Independent Mind-Wandering from EEG using Convolutional Neural Networks"

**Suppl I. Performance of Each Input Type with Different Normalizations.**

The normalization indicates the unit where the normalization is performed:

- *‘Norm Off’* refers to no normalization.
- *‘Norm Channel’* refers to the data from each individual that are normalized within each channel.
- *‘Norm Trial’* refers to the data from each individual normalized within each trial.
- *‘Norm Signal’* refers to the data from each individual is normalized within each time series (i.e., at a channel in a trial).
- ‘*Norm Frequency*’ refers to the data from each individual is normalized by each frequency.
- ‘*Norm Channel x Frequency*’ indicates the data from each individual is normalized per frequency and per channel.
- ‘*Norm Scale*’ indicates the data from each individual is normalized along the scale (one axis of the stERP contour map).
- ‘*Norm Channel x Scale*’ refers to data normalized per scale and per channel.

The normalization is performed using equation: X_norm = (X-min(X))/(max(X)-min(X)). Therefore, the normalized value ranged from 0 to 1.

Bold font indicates the best normalization for each input type according to the performance during validations.

**Raw**

|  | Norm Off | **Norm Channel** | Norm Trial | Norm Signal |
| --- | --- | --- | --- | --- |
| ROCAUC Validation | .495 | **.532** | .500 | .503 |
| ROCAUC Training | .593 | **.670** | .727 | .741 |

**Power**

|  | Norm Off | Norm Channel | Norm Frequency | **Norm Channel x Frequency** | Norm Trial | Norm Signal |
| --- | --- | --- | --- | --- | --- | --- |
| ROCAUC Validation | .503 | .501 | .504 | **.517** | .479 | .493 |
| ROCAUC Training | .534 | 1 | .985 | **.987** | .925 | .971 |

**Inter-Site Phase Clustering (ISPC)**

|  | Norm Off | Norm Channel | Norm Frequency | **Norm Channel x Frequency** | Norm Trial | Norm Signal |
| --- | --- | --- | --- | --- | --- | --- |
| ROCAUC Validation | .493 | .506 | .495 | **.509** | .500 | .498 |
| ROCAUC Training | .823 | .800 | .816 | **.817** | .801 | .806 |

**Single-Trial ERP (stERP)**

|  | Norm Off | Norm Channel | **Norm Scale** | Norm Channel x Scale | Norm Trial | Norm Signal |
| --- | --- | --- | --- | --- | --- | --- |
| ROCAUC Validation | .499 | .484 | **.552** | .533 | .495 | .514 |
| ROCAUC Training | .580 | .673 | **.815** | .815 | .734 | .830 |
