## Supplemental Table 2 for "Decoding Study-Independent Mind-Wandering from EEG using Convolutional Neural Networks"

**Suppl II. Performance of Each Input Type with Different Class Weights**

**Raw**

|  | W_mw_=1 | W_mw_=1.2 | W_mw_=1.4 | **W_mw_=1.6** | W_mw_=1.8 |
| --- | --- | --- | --- | --- | --- |
| ROCAUC Validation | .522 | .507 | .506 | **.530** | .527 |
| ROCAUC Training | .648 | .656 | .657 | **.653** | .668 |

**Power**

|  | W_mw_=1 | **W_mw_=1.2** | W_mw_=1.4 | W_mw_=1.6 | W_mw_=1.8 |
| --- | --- | --- | --- | --- | --- |
| ROCAUC Validation | .499 | **.502** | .492 | .500 | .499 |
| ROCAUC Training | .977 | **.975** | .972 | .972 | .969 |

**Inter-Site Phase Clustering (ISPC)**

|  | W_mw_=1 | W_mw_=1.2 | W_mw_=1.4 | **W_mw_=1.6** | W_mw_=1.8 |
| --- | --- | --- | --- | --- | --- |
| ROCAUC Validation | .488 | .505 | .509 | **.513** | .507 |
| ROCAUC Training | .820 | .811 | .817 | **.810** | .819 |

**Single-Trial ERP (stERP)**

|  | W_mw_=1 | W_mw_=1.2 | **W_mw_=1.4** | W_mw_=1.6 | W_mw_=1.8 |
| --- | --- | --- | --- | --- | --- |
| ROCAUC Validation | .552 | .554 | **.568** | .541 | .548 |
| ROCAUC Training | .812 | .803 | **.804** | .802 | .800 |
