## Supplemental Table 3 for "Decoding Study-Independent Mind-Wandering from EEG using Convolutional Neural Networks"

**Suppl II. Performance of Varying Network Neurons**

|  | [16,16,8] | **[16,32,32]** | [64,64,32] |
| --- | --- | --- | --- |
| ROCAUC Validation | .513 | **.519** | .519 |
| ROCAUC Training | .607 | **.648** | .676 |

*Note* Number in the square brackets denotes the neuron number in CNN layer 1, layer 2 and layer 3.
